# Longitudinal multiomics analysis in blood from individuals before and after HIV acquisition

**DOI:** 10.64898/2026.11.23.767218

**Authors:** Sara Morón-López, Lidia Garrido-Sanz, Leah Soriaga, Víctor Urrea, Emily Wong, Judith Dalmau, Àlex Gómez-Gómez, Noemí Haro, Pep Coll, Beatriz Mothe, Marta Ruiz-Riol, Óscar J. Pozo, Bonaventura Clotet, Christian Brander, Amalio Telenti, Javier Martinez-Picado

## Abstract

A comprehensive understanding of cellular immunity is essential for controlling viral infections, yet individual and coordinated immune responses continue to pose significant challenges in HIV cure research. Recent advances in single-cell and multiomics technologies provide powerful tools to dissect intricate multicellular behaviors and identify novel therapeutic targets. Direct characterization of HIV-induced immune alterations is challenging because matched samples obtained before HIV acquisition are rare. To better understand individual and coordinated cellular immune responses relevant to HIV cure research, we integrated single-nucleus chromatin-accessibility and transcriptomic profiling of 150,060 PBMCs with plasma metabolome, lipidome and soluble immune-profiling from five individuals before HIV acquisition, during early infection and after at least six months of suppressive antiretroviral therapy (ART). HIV acquisition expanded activated CD8 T and proliferating NK/CD8 T cells and induced cell-type-specific chromatin and transcriptional changes enriched for interferon, inflammatory and TNF-NF-κB programs, many persisting during ART. Multiome sequencing detected 32 HIV-positive cells which were associated with increased expression of IFI44L during early infection and ATAD3A after ART. Plasma IP-10 increased transiently, whereas eight energy- or amino-acid-related metabolites showed nominal decreases persisting after ART. These paired observations distinguish transient inflammatory responses from persistent molecular and metabolic perturbations after HIV acquisition.

**Highlights:**

- Longitudinal multiomics single-nuclei data pre- and post-HIV acquisition
- HIV infection disrupts the balance of innate and adaptive immunity
- Cell type-specific molecular alterations: transient and persistent
- Heterogeneous HIV reservoir with distinct epigenomic and transcriptomic states in HIV-positive cells
- Many pathways – particularly metabolism – remain dysregulated after ART initiation

## BACKGROUND

Despite the effectiveness of suppressive antiretroviral therapy (ART), most people living with HIV (PWH) experience viral rebound within a few weeks following ART interruption ^1–5^. This phenomenon underscores that the establishment and persistence of the HIV reservoir remain the main barrier to achieving a cure for HIV. Recent advances in single-cell technologies and multiomic analyses have significantly enhanced our understanding of the virological characteristics of the HIV reservoir and the immune responses associated with HIV infection ^6–11^. However, critical gaps remain in characterizing how pre-acquisition immunity influences HIV reservoir size during both viremic stages and under suppressive ART. Importantly, acquiring matched samples from individuals immediately before and shortly after HIV-1 acquisition is exceptionally rare and logistically challenging, given the unpredictability of infection timing and the need for consistent, longitudinal monitoring.

In this study, we investigated epigenetic and transcriptional changes at the single-nucleus level in peripheral blood mononuclear cells (PBMCs), as well as metabolic and proteomic alterations in plasma, using longitudinal samples from five PWH collected before and during early HIV acquisition (Fiebig stages IV-VI) and after at least six months of suppressive ART, as part of the Check-Ear study cohort (BCN checkpoint). Employing integrated multiomic approaches –including DNA accessibility and gene expression profiling (10x Genomics platform)–we identified immune cell changes following HIV acquisition, chromatin-accessible genes, and transcription factor-binding regions across diverse peripheral immune subsets using assay for transposase-accessible chromatin with sequencing (ATAC-seq), transcriptional alterations via RNA sequencing (RNA-seq), targeted plasma metabolomics, and inflammatory markers through targeted proteomics. This unique access to longitudinal samples before and after HIV-acquisition significantly enhances the relevance and impact of our study, allowing for unprecedented insights into the early immune and molecular changes associated with HIV infection.

Our analysis revealed both transient and persistent global molecular changes that were cell type-dependent, confirmed the cellular heterogeneity of the HIV reservoir, and delineated the impact of HIV infection on the epigenome and transcriptome of HIV positive cells. This study represents a high-dimensional, single-nucleus, longitudinal mapping of epigenetic and transcriptional landscapes in PBMCs, coupled with comprehensive plasma metabolic and proteomic profiling in samples from individuals before and after HIV acquisition and following ART.

## RESULTS

### HIV infection disrupts the balance of innate and adaptive immunity

We utilized integrated single-nucleus ATAC and RNA sequencing (snATAC-seq and snRNA-seq) to analyzed paired PBMCs samples from five individuals collected before and after HIV acquisition (Fiebig stages IV-VI), as well as following at least six months of suppressive ART (**Fig. 1A**). All participants were male, with a median age of 40 years, and were regularly tested for HIV infection and monitored every three months due to high risk of HIV acquisition (Check-Ear and CHAVI cohorts; see **Table 1** for details) ^12^. In total, we profiled 150,060 single-nucleus DNA accessibility and transcriptome profiles from bulk PBMCs, comprising 52,504 nuclei from pre-HIV, 47,868 nuclei from post-HIV, and 49,688 nuclei from post-ART samples across 15 specimens. Sample-specific nuclei and read counts are detailed in **Table S1**. Through high-resolution clustering, we identified 27 immune cell subsets (**Fig. 1B**), including major populations such as CD4, CD8, NK, B, and myeloid cells (**Suppl. Fig. 1**). Comparative analysis of cell proportions across time points revealed a significant expansion of activated CD8 T cells and proliferating NK and CD8 T cells following HIV acquisition, as well as an increased proportion of naïve B cells after suppressive ART (**Fig. 1C**). We also observed a trend toward reduced frequencies of several CD4 T cell-subsets, CD8 T central memory (CM) and terminally differentiated (TEMRA), and B cell subsets and a trend towards an increase in plasmablasts after HIV-acquisition (**Fig. 1C**). These findings demonstrate that HIV infection perturbs the homeostatic balance of both innate and adaptive immune cell subsets.

**Figure 1.**
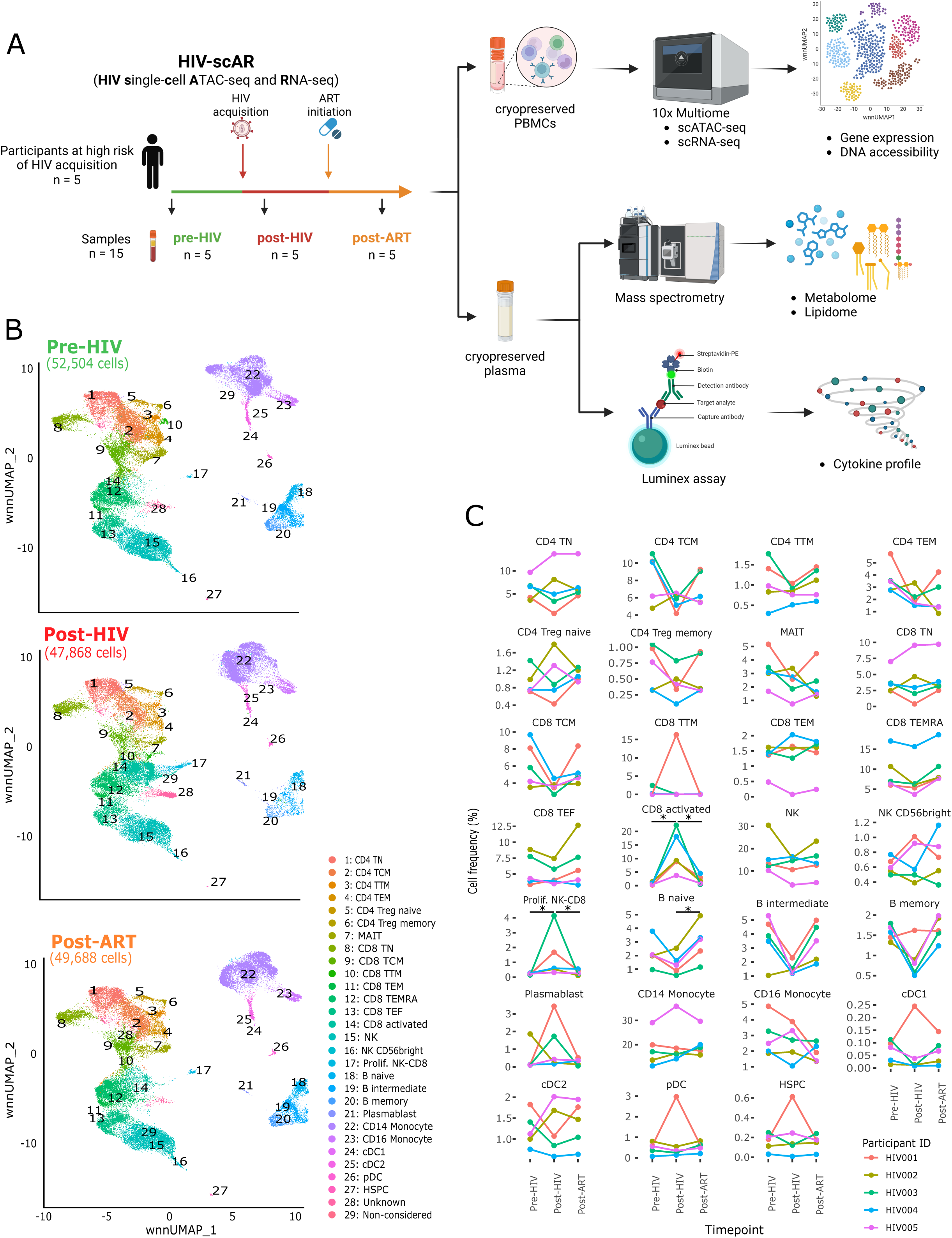
HIV infection disrupts the balance of innate and adaptive immunity. **(A)** Overview of the participants included in this study, the samples collected, and the data generated. Created in BioRender. (B) UMAP visualization shows different immune cell populations identified in pre-HIV acquisition, post-HIV acquisition, and post-ART initiation. The number of captured cells at each time point is indicated in brackets. The T and NK cell compartments include CD4 naïve T cells (CD4 T_N_), CD4 central memory (CD4 T_CM_), CD4 transitional memory (CD4 T_TM_), CD4 effector memory (CD4 T_EM_), CD4 regulatory naïve (CD4 Treg naïve), CD4 regulatory memory (CD4 Treg memory), mucosal-associated invariant T (MAIT), CD8 naïve (CD8 T_N_), CD8 central memory (CD8 T_CM_), CD8 transitional memory (CD8 T_TM_), CD8 effector memory (CD8 T_EM_), CD8 terminally differentiated (CD8 T_EMRA_), CD8 effector (CD8 T_EM_), CD8 activated, natural killer (NK), NK CD56 bright and proliferating NK and CD8 T cells (prolif. NK-CD8). The B cell compartment includes B naïve, B intermediate and B memory cells, and plasmablasts. The myeloid cell compartment includes CD14 monocytes, CD16 monocytes, conventional dendritic cells type 1 (cDC1), conventional dendritic cells type 2 (cDC2), and plasmacytoid dendritic cells (pDC). In addition, we captured hematopoietic stem pluripotent cells (HSPC), an unknown population showing a mix of cell markers and high expression of ribosomal genes, and a non-considered population including a mix of cells with clear identification. These two cell clusters were excluded from further analysis. (C) Individual plots of the dynamics of each cell proportion among the different time points. Each color represents a different participant.

**Table 1.** Clinical data per participant.

| ID | Age at viremia | Sex | Country of origin | HIV subtype | ART at viral suppression | Zenit (copies/ml) | Fiebig<sup>a</sup> | Duration of ART during viral suppression (days) | Nadir (cells/µl) | CD4 count at viremia (cells/µl) | CD4 count at viral suppression (cells/µl) |
|---|---|---|---|---|---|---|---|---|---|---|---|
| 1 | 43 | Male | Spain | B | RAL/TRU | 746,989 | V | 563 | 244 | 244 | 689 |
| 2 | 31 | Male | Spain | B | NVP/TRU | 420,000 | VI | 186 | 366 | 556 | 699 |
| 3 | 58 | Male | Spain | B | DRV/RTV/TRU | 2,600,000 | IV | 425 | 281 | 422 | 537 |
| 4 | 33 | Male | Colombia | B | RAL/TRU | 130,615 | V | 489 | 345 | 403 | 472 |
| 5 | 36 | Male | Argentina | B | GSJ744LA/RPVinjLA | 154,816 | VI | 421 | 847 | 1,207 | 1,766 |

ART, antiretroviral therapy; RAL, raltegravir; TRU, Truvada (emtricitabine and tenofovir); NVP, nevirapine; DRV, darunavir; RTV, ritonavir; GSJ744LA, cabotegravir long-acting; RPVinjLA, rilpivirine long-acting.
<sup>a</sup> Range of days since HIV acquisition by Fiebig stage: IV, 25-30; V, 31-100; VI, 101-183.

### Single-nucleus multiome analysis captured both epigenetic and transcriptional changes of PBMC subsets before and after HIV infection

Examination of snATAC-seq datasets demonstrated that HIV alters DNA accessibility across multiple immune cell types, and this dysregulation is minimally reversed by viremia suppression following ART initiation (**Fig. 2A-C**). Differential accessibility analyses revealed significant gene-level changes when comparing pre-HIV, post-HIV, and post-ART samples in CD4 T naïve (CD4 T_N_), CD4 T central memory (CD4 T_CM_), CD8 T central memory (CD8 T_CM_), CD8 T terminally differentiated (CD8 T_EMRA_), natural killer (NK) and CD14 monocyte cells (**Fig. 2D**, **Table S2 and S3**). Gene set enrichment analysis (GSEA) of these differentially accessible genes highlighted dysregulation of the inflammatory response in NK cells between pre-HIV and post-ART samples, while CD14 monocyte exhibited imbalanced TNF signaling via NF-κB and IFNγ response between pre-HIV and post-HIV samples, and upregulated KRAS signaling, IFNα and IFNγ responses when comparing post-HIV and post-ART samples (**Fig. 2E, Table S3**). Moreover, motif analysis showed altered DNA accessibility in regions associated with specific transcription factors in CD4 T_CM_, CD8 T_CM_, CD8 T_EMRA_, CD8 activated, NK and CD14 monocyte cells across different time points (**Fig. 2F, Table S4**). These findings confirm that HIV acquisition induces either direct epigenetic changes or cell proliferation that evidence the observed changes in DNA accessibility in peripheral immune cells.

**Figure 2.**
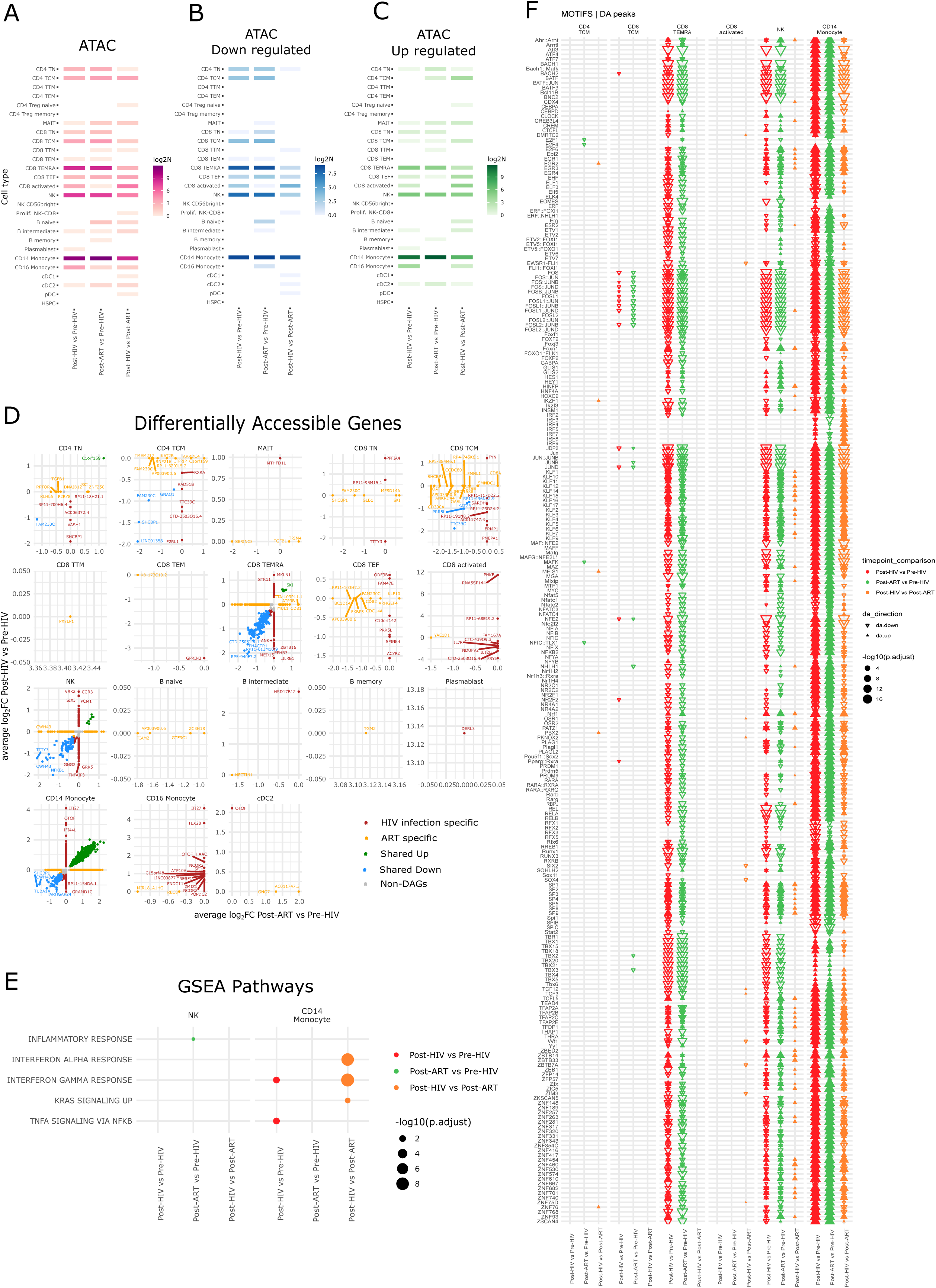
HIV infection dysregulates the epigenetics of PBMCs. (A-C) Number of dysregulated genes per cell type when comparing the different time points (pre-HIV, post-HIV and post-ART), showing the total number of differentially accessible genes (DAGs) associated to peaks (A), the less accessible ones (B) and the more accessible ones (C). (D) Scatter plots of the DAGs per cell type and time point, showing the HIV infection specific ones (red), the ART specific ones (yellow), the shared up (green), the shared down (blue) and the non-DAGs (grey). (E) Significant gene set enrichment pathways detected per cell type when comparing the different time points. (F) Representation of the transcription factors motifs associated with the differentially accessible peaks per cell type and time point comparison.

Analysis of snRNA-seq datasets showed that HIV acquisition impacts gene expression in multiple immune cell types, and in most cases, these changes persist despite viral suppression after ART initiation (**Fig. 3A-C**). Differential expression analysis revealed significant differences across all identified cell clusters except CD8 T transitional memory (CD8 T_TM_), conventional dendritic cells type 1 (cDC1), and hematopoietic stem and progenitor cells (HSPC) (**Fig. 3D**, **Table S5**). The GSEA of differentially expressed genes revealed dysregulation of inflammatory response, IFNα and IFNγ responses, and TNF signaling via NF-κB pathways in nearly all 27 cell clusters (**Fig. 3E, Table S6**), confirming that HIV acquisition alters the transcriptomic landscape of peripheral immune cells.

**Figure 3.**
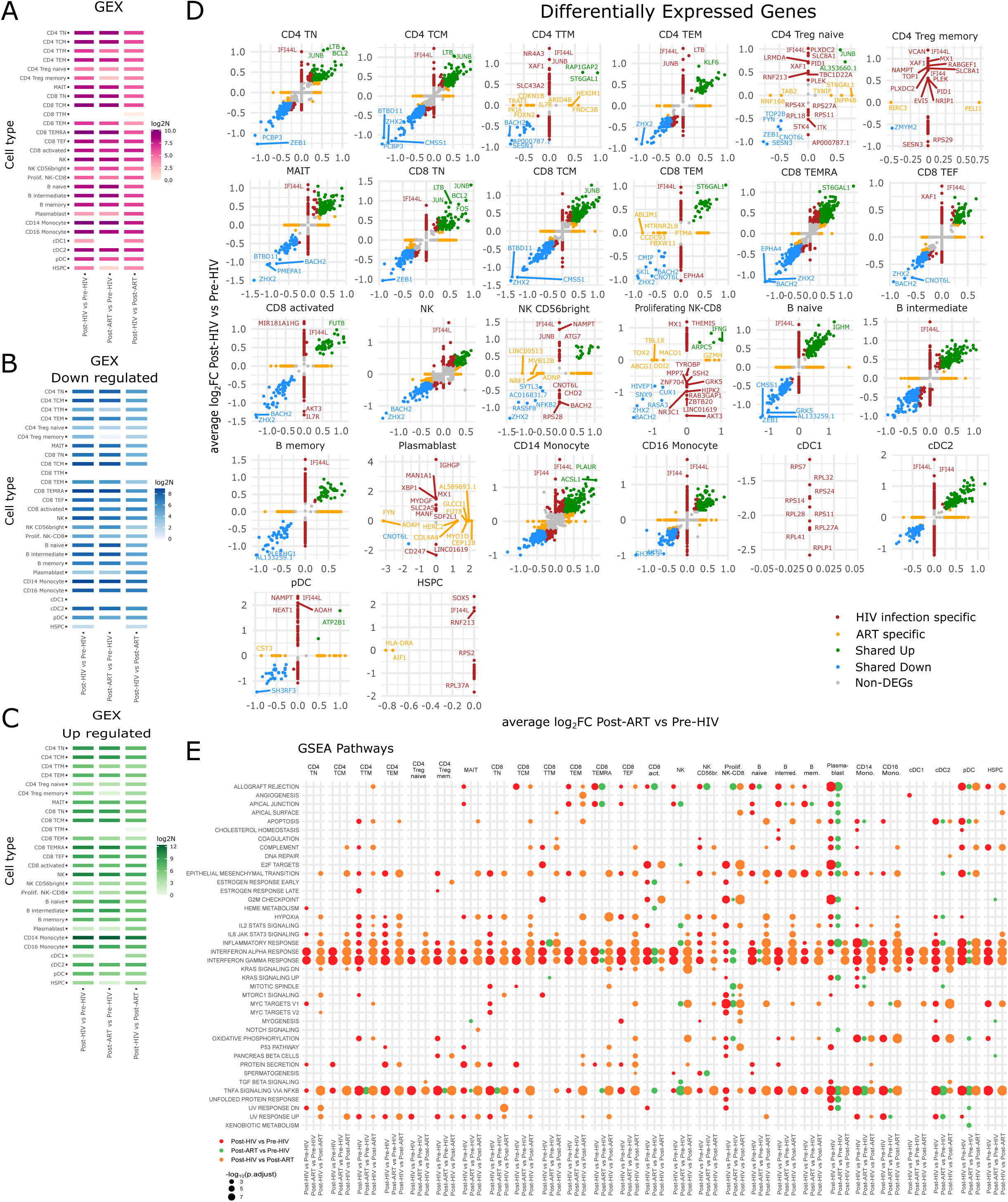
HIV infection dysregulates the gene expression of PBMCs. (A-C) Number of dysregulated genes per cell type when comparing the different time points (pre-HIV, post-HIV and post-ART), showing the total number of differentially expressed genes (DEGs) (A), the down-regulated ones (B) and the up-regulated ones (C). (D) Scatter plots of the DEGs per cell type and time point, showing the HIV infection specific ones (red), the ART specific ones (yellow), the shared up (green), the shared down (blue) and the non-DAGs (grey). (E) Significant gene set enrichment pathways detected per cell type when comparing the different time points.

When integrating the genes with accessible promoters in snATAC-seq and the genes detected in snRNA-seq, we found that 1,985 overlapped between both datasets (**Suppl. Fig. 2A**). Taking into account the levels of DNA accessibility and the gene expression per gene, we found that 850 genes were up-regulated and 410 down-regulated in both datasets, and 725 were discordant between datasets. Overall, we observed that the overlapped genes were predominantly associated with inflammatory response, IFNα, and IFNγ, and TNF signaling via NF-κB (**Suppl. Fig. 2B-C**), consistent with the findings observed in individual analyses.

### Single-cell multiome detection of HIV positive cells reveals genes associated with HIV infection

To confirm HIV infection status and estimate the number of infected cells detectable by single-nucleus analysis, we quantified HIV DNA in PBMCs. As expected, total, intact, 5’defective, and hypermutated/3’defective HIV DNA were negative in pre-HIV samples but were detected in all post-HIV and post-ART samples, with overall levels decreasing after ART initiation (except for intact HIV DNA) (**Fig. 4A**). These results confirm that participants were HIV-negative in pre-HIV samples and that ART effectively reduced HIV reservoir size. The number of HIV-positive cells identified by single-nucleus multiome analysis corresponded with total HIV DNA levels, particularly in post-HIV samples from participants with higher HIV DNA burden, and most notably in the earliest acute infection (Fiebig IV) sample analyzed.

**Figure 4.**
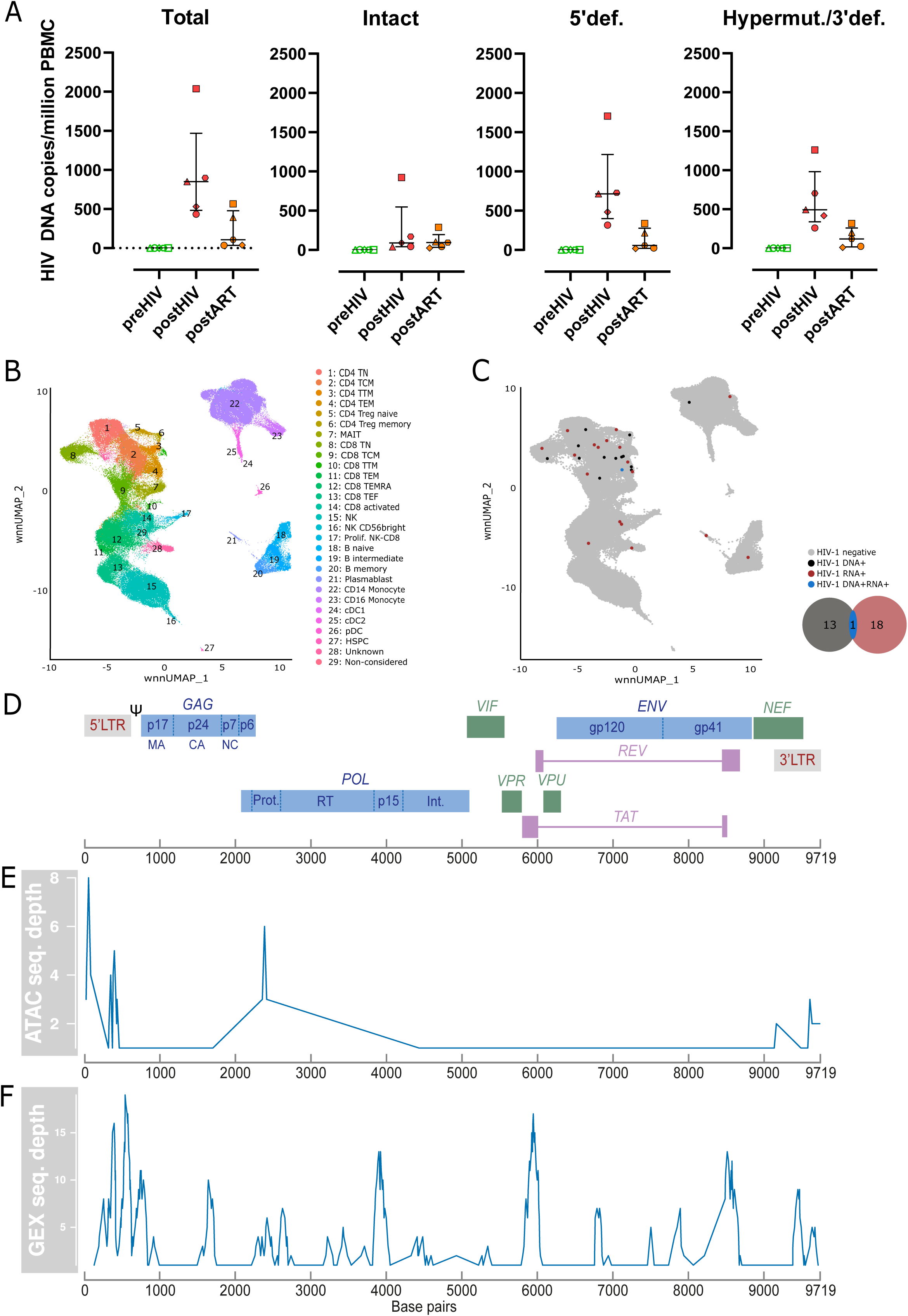
Detection of HIV positive cells. (A) Quantification of total, intact 5’defective (5’def.) and hypermutated/3’defective (hypermut./3’def.) HIV DNA in samples pre-HIV, post-HIV and post-ART from 5 individuals. Median [interquartile range (IQR)] values post-HIV: 850 [482-1,468] copies total HIV DNA/million PBMC; 90 [40-547] copies intact HIV DNA/million PBMC; 714 [399-1,217] copies 5’defective HIV DNA/million PBMC; 494 [339-982] copies hypermutated/3’defective HIV DNA/million PBMC. Median [IQR] values post-ART: 107 [37-478] copies total HIV DNA/million PBMC; 94 [34-196] copies intact HIV DNA/million PBMC; 57 [20-277] copies 5’defective HIV DNA/million PBMC; 119 [18-260] copies hypermutated/3’defective HIV DNA/million PBMC. Median and IQR are represented, and each symbol shape represents a participant. (B) UMAP visualization shows the different immune cell populations identified among all time points. (C) UMAP visualization highlights the HIV positive cells, specifically HIV DNA positive ones in black (n=13), HIV RNA positive ones in red (n=18) and HIV DNA and RNA positive one in blue (n=1). (D) Schematic representation of the HIV-1 genome. (E) Number of HIV DNA reads detected per region of the HIV-1 genome. (F) Number of HIV RNA reads detected per region of the HIV-1 genome.

In total, we identified 32 HIV-positive cells: 26 in the five post-HIV samples and 6 in three of the five post-ART samples. Of these, 13 were detected by snATAC-seq, 18 by snRNA-seq, and 1 cell by both methods. The detection of HIV-positive cells was consistent with HIV DNA quantification in post-HIV and post-ART samples ^13^. Among the HIV-positive cells, 21 were CD4 T cells, 7 were CD8 T cells, 2 were monocytes, and 2 were B cells (**Fig. 4B-C**, **Table S6**). The presence of HIV in cell types other than CD4 T cells may reflect aberrant infection during viremia, active infection of monocytes, or limitations in cell annotation accuracy. Hence, downstream analyses focused exclusively on cells annotated as CD4 T cells based on epigenetic and gene expression profiles, with other cell types excluded from further analyses.

Sequences identified as HIV were aligned to the HIV-1 genome. In DNA accessibility assays, most reads mapped to the LTR regions and *pol* gene, while in gene expression assays, reads were primarily aligned to the 5’LTR (where HIV transcription initiates), as well as *gag* (elongated transcripts), *pol* (unspliced transcripts), and tat-rev and *nef* (multiply-spliced transcripts) (**Fig. 4D-F, Table S7**). These findings confirm that open chromatin regions containing HIV genomes were detected and that active HIV transcription and potential viral protein production were observed, giving that all transcripts were polyadenylated due to the use of single-cell 10x Genomics technology (3’ RNA-seq).

Comparing HIV-positive cells from post-HIV and post-ART samples with uninfected CD4 T cells from pre-HIV samples, we observed significantly higher DNA accessibility and expression of interferon induced protein 44 like (IFI44L) (FDR-adjusted p-value = 0.034, **Fig. 5A-B**) in post-HIV samples, and ATPase family AAA domain containing 3A (ATAD3A) (FDR-adjusted p-value = 0.025, **Fig. 5 C-D**) in post-ART and in joint analysis of HIV-positive cells from post-HIV and post-ART samples. These results are consistent with the observed dysregulation of IFN responses following HIV acquisition (which are normalized after ART initiation) and the persistent metabolic dysregulation after HIV acquisition, as evidenced by DNA accessibility and gene expression analyses.

**Figure 5.**
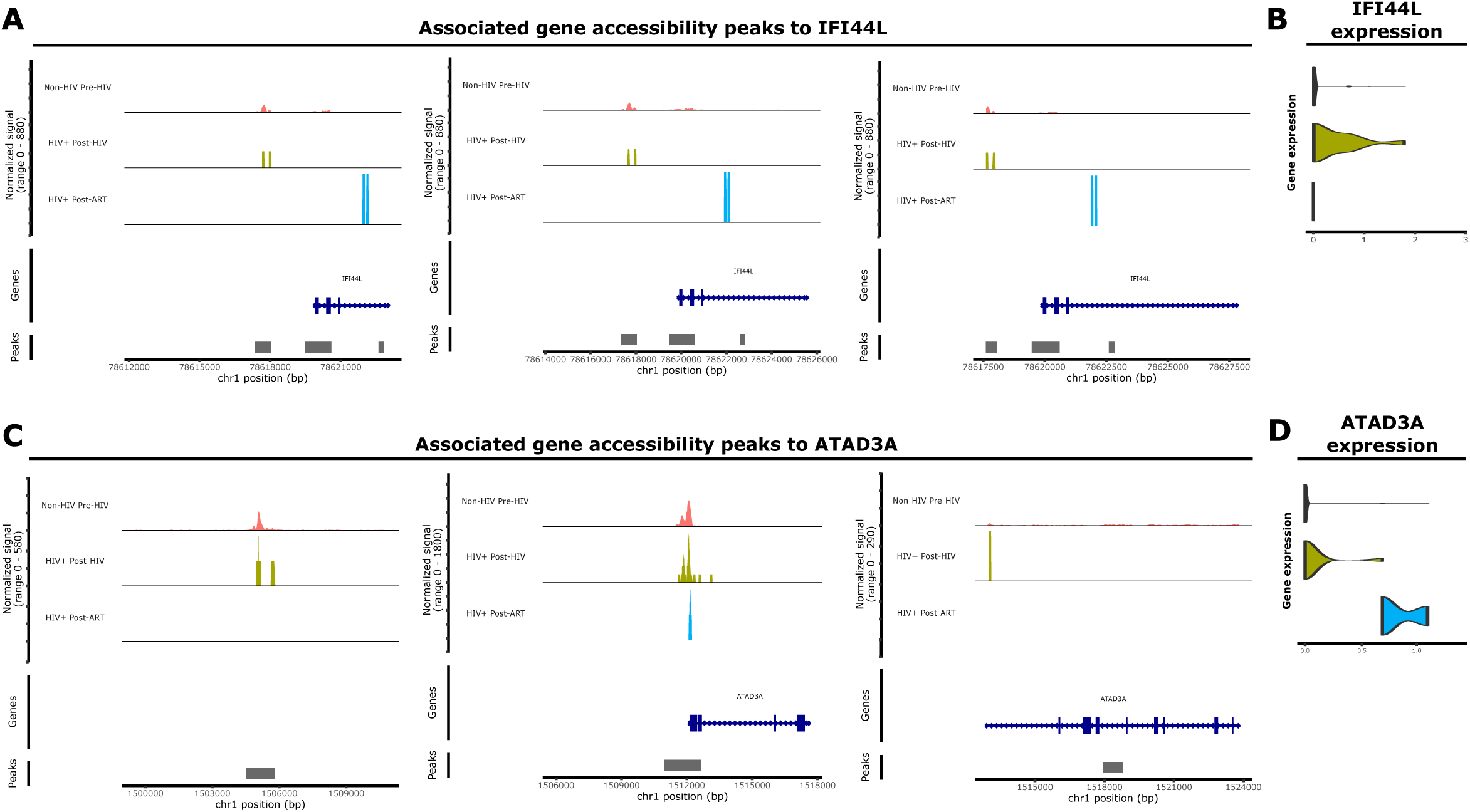
Differential gene accessibility and expression of HIV positive cells compared to HIV negative cells pre-HIV acquisition. (A) Associated gene accessibility peaks to IFI44L per time point. (B) Gene expression of IFI44L per time point. (C) Associated gene accessibility peaks to ATAD3A per time point. (D) Gene expression of ATAD3A per time point.

### HIV infection leads to a transient increase in IP-10 (CXCL10) but persistent alterations in cellular metabolism

To assess changes in plasma inflammatory status following HIV acquisition, we measured 41 cytokines and chemokines. Among these, only IP-10 (CXCL10) showed a significant elevation after HIV infection (p = 0.04, q n.s.) (**Fig. 6A**), which subsequently normalized after ART initiation. We observed an increase in DNA accessibility and gene expression of CXCL10 in CD4 T central memory (CM), CD4 T effector memory (EM), MAIT, CD8 activated, proliferating NK and CD8 T cells, plasmablast and cDC2 cells (**Suppl. Fig. 3A-G**). These findings support previous evidence that HIV infection induces upregulation of IP-10, a chemokine known to facilitate HIV latency and replication^14,15^.

**Figure 6.**
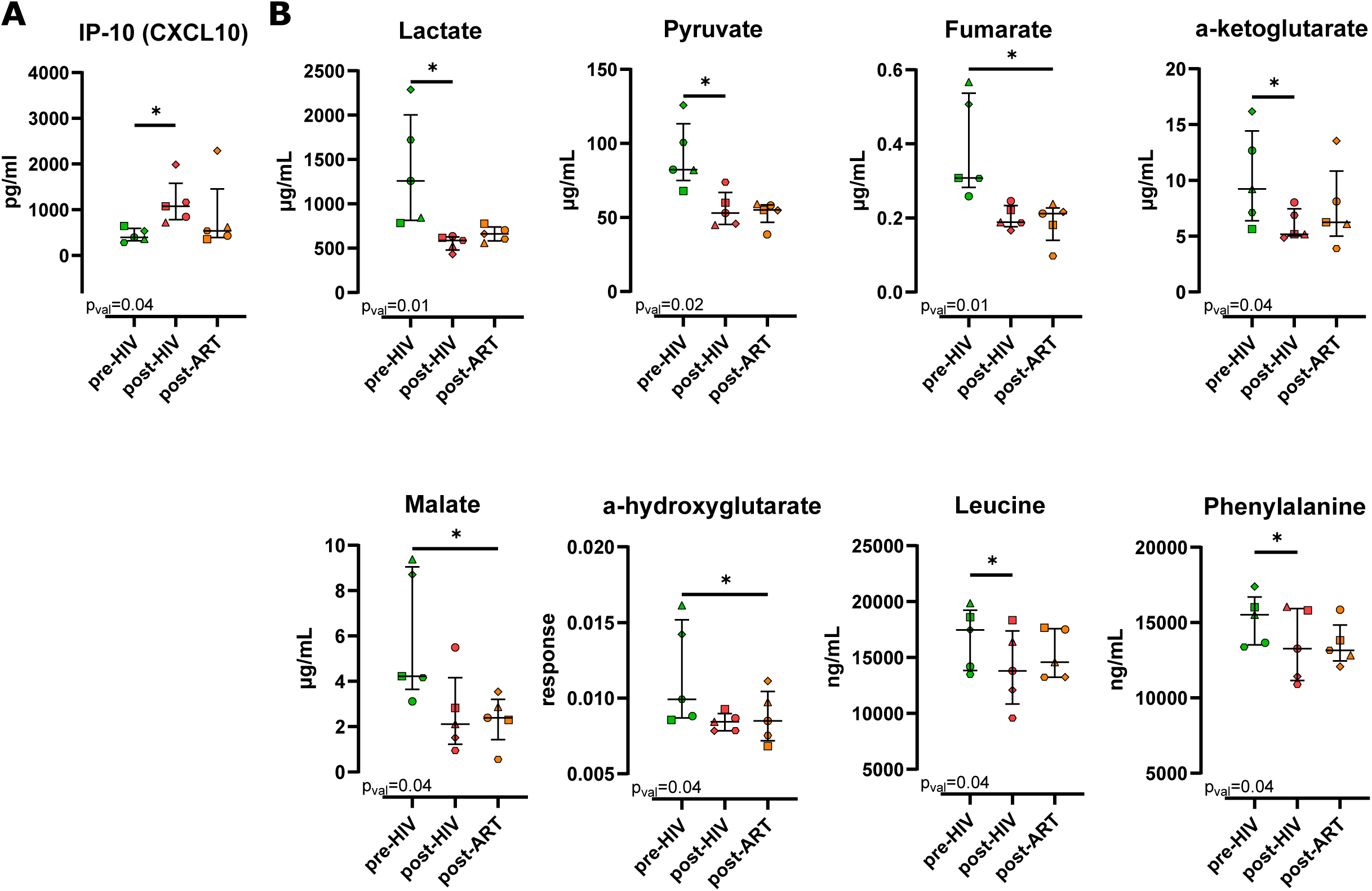
HIV acquisition impacts inflammation markers and metabolites levels in plasma. (A) Levels of IP-10 (CXCL10). (B) Levels of lactate, pyruvate, fumarate, α-ketoglutarate, malate, α-hydroxyglutarate, leucine and phenylalanine in samples pre-HIV, post-HIV and post-ART from the 5 participants. Friedman test and post-hoc Conover test were used to determine the metabolites impacted by HIV infection and ART. Multiple comparison correction was done using false discovery rate (FDR). Median and interquartile range are represented, and each symbol shape represents a participant.

To investigate the impact of HIV infection on plasma metabolome, we quantified 109 metabolites and lipids – based on previous studies –, including 21 associated with energy metabolism, 18 lysophosphatidylcholines, 17 ceramides, 16 related to amino acid metabolism, 15 acylglycerols, 12 compounds linked to microbiota, 5 bile acids, and 5 choline metabolism-related compounds. Of the metabolites and lipids analyzed, six metabolites involved in anaerobic and aerobic metabolic pathways–specifically the citric acid cycle (lactate, pyruvate, α-ketoglutarate, fumarate, malate and α-hydroxyglutarate)–as well as two amino acids (leucine and phenylalanine), which contribute to Acetyl-CoA production, were significantly decreased after HIV infection (all p<0.05, q n.s.). Notably, these reductions persisted and were not restored following at least six months of suppressive ART (**Fig. 6B**). These results indicate that HIV infection disrupts cellular metabolism, and that these metabolic perturbations are not reversed by short-term ART.

## DISCUSSION

In this study, we analyzed blood samples collected from the same individuals before and after HIV acquisition, as well as during ART-mediated viral suppression. This longitudinal approach enabled us to assess how HIV infection and subsequent ART initiation affect epigenetic and transcriptional responses across peripheral immune cells, and how this data correlates with metabolomic, lipidomic and proteomic plasma markers. We performed single-nucleus epigenetic and transcriptomic profiling of a total of 150,060 PBMCs: 52,504 pre-HIV acquisition, 47,868 post-HIV acquisition, and 49,688 post-ART initiation (at least 6 months after confirmed viral suppression).

Among the single nuclei analyzed, using our own developed bioinformatic pipeline to detect HIV sequences, we identified 32 HIV-positive cells: 26 in post-HIV samples and 6 in post-ART. Of these, 13 were identified in snATAC-seq data, 18 in snRNA-seq data, and 1 cell in both datasets. Despite the relatively low number of HIV-positive cells, we detected HIV genomic reads aligning to the 5’ LTR, the TAR loop, the initial region of *pol* and the 3’LTR. These regions likely correspond to open chromatin at the HIV promoter, and HIV transcript reads spanned the entire viral genome, with enrichment in the TAR loop and tat regions, suggesting active HIV transcription. This observation is consistent with previous findings using DOGMA-seq ^6^ and HIV-seq ^9^. Furthermore, the number of HIV-positive cells identified corresponded with HIV DNA levels quantified in PBMCs, supporting the reliability of our multi-reference, Kraken-2-based bioinformatic pipeline ^13^.

Our results demonstrate dysregulation of both innate and adaptive immune responses following HIV acquisition and during suppressive ART, characterized by significant increases in activated CD8 T cells and proliferating NK and CD8 T cells. We confirmed that HIV acquisition alters the epigenetic and transcriptomic landscape of peripheral immune cells, with changes primarily associated with inflammatory responses, IFNα, and IFNγ, and TNF signaling via NF-κB. These findings are consistent with previous studies that utilized single-cell RNA-sequencing in PBMCs from untreated females living with HIV before and during acute infection^8^ as well as single-cell/nuclei multiome in PBMCs from untreated individuals living with HIV ^16^. Notably, proliferating NK cells have been implicated in HIV control ^8^, and enhanced NK cell function and activation are associated with reduction in HIV reservoir size ^17–19^.

Focusing on HIV-positive CD4 T cells and comparing them to HIV-negative CD4 T cells from pre-HIV samples, we identified upregulation of IFI44L in post-HIV samples and ATAD3A in post-ART samples. IFI44L is an IFN-stimulated gene implicated in innate and antiviral immune responses, with its expression modulated by both IFN signaling and HIV infection ^20,21^. Rather than functioning as a direct viral inhibitor, IFI44L acts as a regulator that mitigates inflammation, thereby protecting the host from chronic overactivation of the immune system ^20–23^. The observed upregulation of IFI44L following HIV acquisition likely reflects increased inflammation induced by the infection, which is subsequently normalized after ART initiation. In contrast, ATAD3A encodes a mitochondrial membrane protein essential for maintaining cellular metabolism, mitochondrial integrity, and nucleoid organization. During viral infections, ATAD3A can interact with viral proteins and modulate innate immune responses ^24,25^. The sustained upregulation of ATAD3A after HIV acquisition, which does not revert to pre-HIV levels following ART, may be indicative of persistent metabolic dysregulation in infected cells. These findings mirror alterations in plasma immune and metabolic markers following HIV-acquisition, with metabolic disturbances persisting despite ART-mediated viral suppression. Specifically, we observed transient increases in plasma IP-10 (CXCL10) after HIV acquisition that normalized following ART, sustained reductions in multiple metabolites related to the Krebs cycle, which persisted even after ART. These results align with previous research in inflammatory and metabolic changes in PWH before and after ART initiation ^26–30^. Nonetheless, this study provides valuable multiomics data from human samples both before and after HIV acquisition, demonstrating that metabolic dysregulation arises after HIV acquisition and is not restored after at least six months of effective ART.

Several limitations should be acknowledged. Firstly, the low number of HIV-infected cells identified during viremia and viral suppression may constrain the conclusions drawn from comparisons between HIV-negative and HIV-positive cells. However, the use of longitudinal pre-HIV acquisition samples from the same individuals may reduce inter-individual variability and reinforce observed differences. Second, limited HIV DNA detection by ATAC-seq, with more than half of the HIV-positive cells classified as HIV DNA-negative but RNA-positive, highlights the lower resolution of single-cell ATAC-seq compared to plate-based methods ^31^, which may increase HIV-positive read recovery but reduce throughput for clinical samples. Third, our study was performed in single nuclei rather than whole cells, and participants were young males from Spain and South America infected with HIV subtype B ^12^. Thus, our findings may not be generalized to studies employing single cell approaches, or to populations of different ages, sexes, ethnicities or HIV subtypes. However, some results are consistent with those from the FRESH study, which used single-cell analysis of PBMCs from females before and after acute HIV acquisition ^8^, suggesting that at least some conclusions, such as the increase in cytotoxic cell subsets and the up-regulation of interferon-stimulated genes, may be applicable across methodologies, sexes and HIV subtypes.

Overall, our study advances understanding of how HIV infection impacts peripheral blood cellular dynamics and immune-metabolism, highlighting the potential role of proliferating NK and CD8 T cells in controlling HIV persistence. Therapeutic strategies aimed at restoring pre-infection metabolic states and enhancing the activity of proliferating NK and CD8 T cells may improve the HIV control and warrant further investigation as potential approaches toward an HIV cure.

## CONCLUSIONS

Overall, this work establishes a longitudinal, single-nucleus multi-omic framework for dissecting the cellular and molecular consequences of HIV acquisition and treatment. By revealing extensive cell type-specific epigenetic and transcriptional remodeling, persistent immunometabolic perturbations, and the pronounced heterogeneity of the HIV reservoir, our study provides a comprehensive view of the biological imprint left by HIV infection. Importantly, the identification of proliferating NK and CD8 T cell populations as potential regulators of viral persistence, together with the association between HIV control and restoration of pre-infection metabolic states, uncovers actionable mechanisms that could be leveraged therapeutically. These findings refine our understanding of HIV pathogenesis and nominate immune and metabolic reprogramming as complementary targets for strategies aimed at achieving durable HIV remission and ultimately a cure.

## METHODS

### Study participants

Participants were recruited at the BCN Checkpoint (Barcelona, Spain; Check-Ear cohort) and followed up at Infectious Diseases Foundation and Immunopathologies Research Institute IrsiCaixa (Badalona, Spain; CHAVI cohort) after being diagnosed with HIV infection. The influence of gender was not measured in this study, as all the participants were cisgender men. Peripheral blood mononuclear cells (PBMCs) were obtained according to protocols approved by the Institutional Review Board of Germans Trias i Pujol Hospital. Clinical characteristics of study participants and samples are summarized in Table 1.

### Study approval

Study participants were tested for HIV infection at the BCN Checkpoint facilities and the HIV unit of Barcelona’s Clinic hospital and followed by Germans Trias i Pujol Hospital. Participants gave written informed consent to participate in accordance with the Declaration of Helsinki. The study was approved by the local independent institutional review boards (Check-Ear: EO-09-014, and CHAVI: EO-08-023).

### Sample processing and PBMC isolation

Blood samples were collected in EDTA tubs and immediately processed. Plasma was collected after centrifugation and stored at -80°C. PBMCs were isolated using ficoll plaque, cryopreserved in FBS with 10% DMSO, stored for overnight at -80°C and then transferred to liquid nitrogen.

### Single-nuclei 10x Multiome ATAC-Seq and Gene Expression data generation

#### Sample preparation

Cryopreserved PBMCs were rapidly thawed in a 37°C water bath. Each sample was transferred into a 15 ml Falcon with RPMI supplemented with 50% FBS and immediately centrifuge at 300xg for 5 min. The supernatant was removed without disrupting the cell pellet, resuspended in 1 ml PBS + 0.04% BSA, filtered using a 40 µm Cell Strainer and transferred to a new 15 ml Falcon. Cell count and viability were assessed with LUNA automatic counter (Cell viability >85% for all samples).

Nuclei isolation was performed following the “Nuclei Isolation for Single Cell Multiome ATAC + Gene expression sequencing” demonstrated protocol (10x Genomics, CG000365 Rev. C). Briefly, 1×10^6^ PBMC were transferred to a 1.5 ml microcentrifuge tube and centrifuged at 300xg for 5 min at 4°C. The supernatant was removed without disturbing the cell pellet, and 100 µl of chilled Lysis Buffer (10mM Tris-HCl (pH 7.4); 10 mM NaCl; 3 mM MgCl_2_; 0.1% Tween-20; 0.1% Nonidet P40 Substitute; 0.01% Digitonin; 1% BSA; 1 mM DTT; and 1U/µl RNase inhibitor) was added and pipette-mixed 10 times. Samples were then incubated on ice for 3 min. Following the lysis, 1 ml of chilled Wash Buffer (10 mM Tris-HCl (pH 7.4); 10 mM NaCl; 3 mM MgCl_2_; 1% BSA, 0.1% Tween-20; 1 mM DTT; and 1 U/µl RNase inhibitor) was added and pipette-mixed 5 times. Nuclei were centrifuged at 500xg for 5 min at 4°C, and the supernatant removed without disrupting the nuclei pellet. Based on the cell counting and assuming a 50% nuclei loss during cell lysis, nuclei were resuspended into the appropriate volume of chilled Diluted Nuclei Buffer (1X Nuclei Buffer (20X, 10x Genomics Single Cell Multiome ATAC Kit A); 1 mM DTT; and 1 U/µl RNase inhibitor) in order to achieve a nuclei concentration of 3,230-8,060 nuclei/µl, suitable for a Target Nuclei Recovery of 10,000. The resulting nuclei concentration was determined with LUNA automatic counter and samples were immediately processed following the “Chromium Next GEM Single Cell Multiome ATAC + Gene Expression” user guide (10x Genomics, CG000338 Rev. E) to prepare the single-cell ATAC-seq (scATAC-seq) and Gene Expression (GEX) libraries. Briefly, the transposition reaction was prepared by pipette-mixing the desired number of nuclei, following Nuclei Concentration Guidelines, with the Transposition Mix (70% ATAC Buffer B + 30% ATAC Enzyme B; 10x Genomics), and incubated in a thermal cycler at 37°C for 60 min. Then, immediately, the Master Mix (82.5% Barcoding Reagent Mix + 1.8% Template Switch Oligo + 3.2% Reducing Agent B + 12.5% Barcoding Enzyme Mix; 10x Genomics) was pipette-mixed with the transposition reaction and loaded into a Chromium Next GEM Chip J, followed by adding the Single Cell Multiome Gel Beads (10x Genomics) and the Partitioning Oil (10x Genomics). GEMs were generated using a Chromium Controller and immediately transferred into tube strips and incubated at 37°C for 45 min followed by 25°C for 30 min. After this cycle, Quenching Agent (10x Genomics) was added to each sample and GEM cleanup was performed using Dynabeads MyOne SILANE (10x Genomics) and SPRIselect reagent (Beckman Coulter). Then, Pre-Amplification Mix (92.6% Pre-Amp Mix + 7.4% Pre-Amp Primers; 10x Genomics) was added to each sample and incubated using the following thermal cycling conditions: 72°C for 5 min, 98°C for 3 min, 7 cycles of 98°C for 20 seconds, 63°C for 30 seconds and 72°C for 1 min, and a final extension of 72°C for 1 min. Pre-Amplification PCR products were cleaned up using SPRIselect reagent (Beckman Coulter). SPRI cleaned sample included barcoded transposed DNA fragments and barcoded cDNA. This mix was divided and used as input for ATAC Library Construction and for cDNA Amplification, which was used for Gene Expression Library Construction.

##### Library preparation

ATAC Library Construction was performed by adding the Sample Index PCR Mix (87% Amp Mix + 13% SI-PCR Primer B; 10x Genomics) to the pre-amplified sample, followed by the addition of individuals Sample Index N, Set A (10x Genomics), and incubation using the following thermal cycle conditions: 98°C for 45 seconds, 7 cycles of 98°C for 20 seconds, 67°C for 30 seconds and 72°C for 20 seconds, and a final extension of 72°C for 1 min. Then, post sample Index Double Sided Size Selection was done using SPRIselect reagent 0.6X (Beckman Coulter) and post library construction quality was verified using an Agilent Bioanalyzer High Sensitivity chip (average peak range size: 373-517 bp).

cDNA Amplification was performed by adding cDNA Amplification Mix (77% Amp Mix + 23% cDNA Primers; 10x Genomics) to the pre-amplified sample, and incubating using the following thermal cycle conditions: 98°C for 3 min, 6 cycles of 98°C for 15 seconds, 63°C for 20 seconds and 72°C for 1 min, and a final extension of 72°C for 1 min. Later, amplified cDNA was cleaned up using SPRIselect reagent (Beckman Coulter) and the quality and quantification was assessed on an Agilent Bioanalyzer High Sensitivity chip. Next Gene Expression Library Construction was done by adding Buffer EB (10x Genomics) and Fragmentation Mix (33% Fragmentation Buffer + 67% Fragmentation Enzyme; 10x Genomics) to the amplified cDNA, followed by incubation in a pre-cooled thermal cycler at 4°C following the thermal conditions: 32°C for 5 min and 65°C for 30 min. Then the samples were cleaned up using SPRIselect 0.6X (Beckman Coulter) and the adaptor ligation was done by adding the Adaptor Ligation Mix (40% Ligation Buffer + 20% DNA Ligase + 40% Adaptor Oligos; 10x Genomics) and incubated in a thermal cycler at 20°C for 15 min. Post Ligation samples were cleaned up using SPRIselect 0.8X (Beckman Coulter) and Index PCR was performed by adding the Amp Mix and the individual Dual Index TT Set A under the following thermal cycler conditions: 98°C for 45 seconds, 13-15 cycles (depending on cDNA input) of 98°C for 20 seconds, 54°C for 30 seconds and 72°C for 20 seconds, and a final step of 72°C for 1 min. PCR products were cleaned up using SPRIselect 0.6X (Beckman Coulter) and quality of the library was assessed on an Agilent Bioanalyzer High Sensitivity chip (average peak range size: 452-505 bp).

##### Sequencing

Finally, libraries were loaded on an Illumina NovaSeq 6000 system. Chromium Single Cell Multiome ATAC libraries comprise double stranded DNA with standard Illumina paired-end constructs which begin with P5 and end with P7. Sequencing these libraries produces a standard Illumina BCL data output folder that includes paired-end Read 1N and Read 2N used for sequencing the DNA insert, 8 bp sample index in the i7 read, and 24 bp in the i5 read to cover the 16 bp 10x Barcode and 8 bp Spacer. Chromium Single Cell Multiome Gene Expression libraries comprise cDNA insert with standard Illumina paired-end constructs which begin with P5 and end with P7. Sequencing these libraries produces a standard Illumina BCL data output folder. TruSeq Read 1 is used to sequence 16 bp 10x Barcodes and 12 bp UMI, while 10 bp i5 and i7 sample index sequences are the sample index reads. TruSeq Read 2 is used to sequence the insert. FASTQ files were obtained from the BCL data using the blc2fastq2 v2.20 software.

### Droplet digital PCRs

Pellets of 1×10^6^ PBMCs were collected after thawing, resuspended in Lysis buffer consisted of 0.1% Triton x-100 (Sigma) and 400 μg/ml Proteinase K (Ambion) in 10 mM Tris-HCl pH = 9.0 (Sigma). Cell lysates were incubated at 55°C for 15-20h, followed by a 5 min inactivation of proteinase at 95°C, prior to storage at -20°C. To quantify total HIV DNA, we used two different primers/probe sets annealing to the 5’LTR and GAG conserved regions of HIV-1 genome to circumvent potential primer mismatch in individuals’ viral sequence ^32–34^. To quantify intact, 5’defective and hypermutated/3’defective HIV DNA we used primers/probes sets annealing to the Packaging signal and Env Rev Response Element regions following the intact proviral DNA assay (IPDA) protocol ^35^. ddPCR Supermix for Residual DNA quantification was used (186-4037, Bio-Rad). PCR was done using the C1000 Touch™ Thermal Cycler (Bio-Rad) and subsequently analyzed using a QX100™ droplet reader (Bio-Rad) and the QuantaSoft v.1.6 software (Bio-Rad) ^36^. Samples were quantified in duplicate. PBMCs from HIV-negative donors were used as negative controls and assayed in each plate to set the positive/negative threshold for ddPCR analysis, and the number of those negative control wells was the same as replicas for each sample. The RPP30 cellular gene was quantified in parallel to normalize sample input and assess DNA shearing.

### Targeted plasma metabolomics and lipidomics

After protein precipitation, plasma samples were analyzed via five protocols, which assessed each family of compounds using previously reported methods ^37–40^. A set of 109 targeted metabolites belonging to preselected metabolic pathways was determined by selected reaction monitoring by LC–MS/MS system consisting of an Acquity UPLC instrument (Waters Associates, Milford, MA, USA) coupled to a triple quadrupole (TQS Micro, Waters) mass spectrometer. The metabolite panel contained 16 compounds related to amino acid metabolism, 21 related to energy metabolism, 15 acylglycerols, 17 ceramides, 18 lysophosphatidylcholines (LPCs), 12 compounds related to microbiota, 5 biliary acids, and 5 compounds of choline metabolism. MassLynx software V4.1 (Waters Associates) was used for peak integration and data management.

### Cytokine determination

After thawing at 37°C, 41 cytokines/chemokines (EGF, Eotaxin, G-CSF, GM-CSF, IFNα2, IFNγ, IL-10, IL-12P40, IL-12P70, IL-13, IL-15, IL-17A, IL-1RA, IL-1α, IL-1β, IL-2, IL-3, IL-4, IL-5, IL-6, IL-7, IL-8, IP-10, MCP-1, MIP-1α, MIP-1β, RANTES, TNFα, TNFβ, VEGF,FGF-2, TGF-α, FIT-3L, Fractalkine, GRO, MCP-3, MDC, PDGF-AA, PDGF-AB/BB, sCD40L, and IL-9) were quantified using a MILLIPLEX® Map Human Cytokine/Chemokine Magnetic Bead Panel (Luminex®, HCYTMAG-60K-PX41), following the manufacturer’s instructions. Briefly, each sample, controls and standards were quantified in duplicate. Neat plasma was used to quantify 38 cytokines using the premixed beads of the kit, while 1/100 diluted plasma was used to measure RANTES, PDGF-AA and PDGF-AB/BB using the specific mix of bead, as recommended by the manufacturer’s instructions. Plates were incubated 2 h at RT on a plate shaker (500-800 rpm); then, washed, detection antibodies added and incubated with agitation on a plate shaker for 1 h at RT (500-800 rpm). Then, Streptavidin-Phycoerythrin was added to each well, and the plates incubated on a plate shaker for 30 min at RT (500-800 rpm). After this incubation, plates were washed, sheath fluid added, and beads resuspended on a plate shaker for 5 min. The plates were read immediately using Luminex® 200 (Luminex Corp).

### Bioinformatics and statistics

Raw multiome FASTQ reads (snATAC-seq and snRNA-seq) were quality control tested with FASTQC (v0.11.9) ^41^. Reads were aligned to the GRCh38 reference genome provided by 10x Genomics (refdata-cellranger-arc-GRCh38-2020-A-2.0.0.tar.gz) using cellranger-arc count (10x Genomics v2.0.2). All downstream analysis and graphical representation were performed with R (v4.3.0) ^42^ on a controlled environment generated with renv package (v0.17.3).

Feature matrices were cleaned from background contamination, and empty droplets were removed by SoupX package (v1.6.2) ^43^. Count matrices were used to create a Seurat object using EnsDb.Hsapiens.v86 package (v2.99.0) ^44^ for chromatin annotation based on Ensembl repository. Then, to ensure the high-quality of the data, samples were processed with Seurat (v4.3.0) ^45^ and Signac (v1.9.0) ^46^ packages to remove cells from the analysis if they met any of the following criteria: (1) expressing few genes; (2) having too high or low amount of UMI counts and ATAC molecules; (1) and (2) thresholds were set based on 1.5 IQR; (3) TSS enrichment above 1 and a nucleosome signal below 2 ^47^; (4) mitochondrial percentage above 20%; and, (5) likely doublets identified using scDblFinder package (v1.14.0) ^48^. Mitochondrial genes were eliminated from the matrix.

UMI counts were normalized using a sctranform-based approach and a Principal Component Analysis (PCA) was run to reduce the dimensionality of the data. Individual samples were integrated based on selected features and using reciprocal PCA (RPCA) on a reference-based approach. The resulting object was again normalized as before and performed a linear and non-linear dimensional reduction of the data with PCA and the Uniform Manifold Approximation and Projection (UMAP), respectively. Nearest neighbours were identified in the mutual nearest neighbour space and were used to define clusters with resolution argument set to 0.5. On snATAC-seq data, peaks were identified for each RNA defined cluster using MACS2 ^49^, and peaks located either in non-standard or in genomic blacklist were removed. For each peak, fragment counts were quantified and normalized with term-frequency inverse-document-frequency (TF-IDF) method ^50^. The new matrix was used for UMAP representation generated with latent semantic indexing (LSI) reduction.

snRNA-seq and snATAC-seq data were integrated using the Weighted Nearest Neighbor (WNN) graph and visualized with UMAP. Cells were clustered using a resolution of 0.8. Individual cells were annotated using human PBMC reference from Azimuth app ^47,51^ and VNPs reference ^52^. We further manually curated the annotation of clusters using information from prediction scores and marker genes characteristic from specific cell type. Next, differential abundance of cell types across time points was analysed using log2 fold-change of relative proportions.

Cell type specific markers and differentially expressed genes (DEGs) between timepoints were identified. We quantified the magnitude of the gene expression changes across timepoints by totalling the number of DEG with an adjusted p-value < 0.05 and absolute log2 fold change > 0.25). Next, peaks were linked to genes and their closest feature. For each cell type, we also identified the genes associated to differentially accessible peaks (DAPs) between timepoints. The magnitude of the changes on peaks accessibility was quantified counting the total number of differentially accessible peaks with an adjusted p-value < 0.1 and absolute log2 fold change > 0.25.

Gene Set Enrichment Analysis (GSEA) was run for both DEGs and on genes associated to DAPs linked to genes using clusterProfiler package’s (v4.8.1) ^53,54^ using the annotation from the H collection-hallmark gene sets from msigdbr package (v7.5.1) ^55^. P-values from enriched gene sets were adjusted using “fdr” method and were filtered using an adjusted p-value cut-off of 0.05 and 0.10 for DEGs and DAPs, respectively. For enrichment analysis of genes associated with peaks, only DAPs with a p-value <=0.10 and absolute log2 fold change >0.25 were selected. Motif sequences were also identified with Signac.

HIV-1 infected cells were determined using a custom pipeline^13^. Briefly, we developed an in-house pipeline to identify infected cells based on Kraken2 ^56^ classification of reads and custom virKra R package (v1.0.0) (https://github.com/irsi-grec/virKra) to extract cell-specific barcodes. Multiome reads were processed to determine DNA fragment and UMI counts associated with HIV by classifying reads to the virus with Kraken2 and a reference database populated with 1,312 HIV genome sequences from Los Alamos HIV Sequence Database (accessed on 2022-12-19). HIV associated reads were transformed into counts and transferred into the Seurat object with virKra. To find genetic markers associated with the HIV-1 infection, CD4 T cells from pre-HIV samples were compared against HIV positive CD4 T cells from post-HIV and post-ART samples, separately and infected samples together. The coverage of viral sequences was generated by aligning HIV-1 reads to two different HIV-1 subtype B references using BWA (v0.7.17-r1188) with MEM algorithm ^57^; snATAC-seq reads were mapped against AF256204 genome and snRNA-seq reads against X01762 genome. Resulting SAM files were converted to BAM format with samtools (v1.10) ^58^. Genome coverages were recovered using Bedtools (v2.29.2) ^59^ and plotted with Gviz (v1.44.2) package ^60^.

Metabolites, lipids, cytokines, and chemokines levels were analyzed using Friedman test (p-value) and post-hoc Conover test (p-value between timepoints) and corrected by multiple comparison using FDR (q-value).

## DECLARATIONS

### Ethics approval and consent to participate

Participants gave written informed consent to participate in accordance with the Declaration of Helsinki. The study was approved by the local independent institutional review boards at Barcelona’s Clinic hospital and Germans Trias i Pujol Hospital (Check-Ear: EO-09-014, and CHAVI: EO-08-023).

### Consent for publication

JM-P as co-corresponding author confirms that all authors have read and agreed with the current version of the manuscript.

### Availability of data and materials

#### Lead contact

Further information and requests for resources and reagents should be directed to and will be fulfilled by the lead contacts, Javier Martinez-Picado and Sara Morón-López.

#### Materials availability

This study did not generate new unique reagents.

#### Data and code availability

- snATAC-seq data was deposited to European Genome-phenome Archive (EGA) with accession number available upon acceptance.

- snRNA-seq data was deposited to Genome-phenome Archive (EGA) with accession number available upon acceptance.

- Metabolome, lipidome, HIV-1 DNA and immune-soluble markers metadata was deposited to Consortium of University Services of Catalonia (CSUC) with accession number with accession number available upon acceptance.

- This paper reports specific and original code available on GitHub upon acceptance.

- Any additional information required to reanalyze the data reported in this paper is available from the lead contact upon request.

### Competing interests

The authors declare no competing interests. J.M-P. has received institutional grants and educational/consultancy fees from Grifols, Merck Sharp & Dohme and ViiV Healthcare, all outside the submitted work. S.M-L. has received institutional grants from Gilead Sciences outside the submitted work. PC has received institutional grants and educational/consultancy fees from Merck Sharp & Dohme, Gilead Sciences and ViiV Healthcare, all outside the submitted work.

### Funding

SM-L was supported by the EU Gilead Research Scholars Program in HIV award 2021, grant 2020 BP 00046 from the Catalan Agency of Management of University and Research Grants and grants PID2023-150316OA-I00 and RYC2024-048369-I from the Spanish Ministry of Science and Innovation. LG-S received financial support by the grant SLT02823 000257, funded by the "Strategic plan for research and innovation in health" (PERIS), from the Catalan Department of Health. CB, MR-R, BM were supported by European Commission Horizon Europe GA6932308. “Epigenetic regulation of host factors in viral infections” (EPIVINF). Research in JM-P’s laboratory related to this project is supported by the Spanish Ministry of Science and Innovation (grant PID2022-139271OB-I00), CIBERINFEC (CB21/13/00063, Spain), and NIH/NIAID (1 UM1 AI164561-01 and 1P01AI178376-01, United States of America). Sample processing at IrsiCaixa was possible thanks to the crowdfunding initiative of People in Red and Grifols S.A.. We thank CERCA Programme/Generalitat de Catalunya for institutional support (2025 SGR 01633).

## Acknowledgments

Firstly, we thank all the participants and BCN Checkpoint, without whom this study could not have been performed. We are grateful to the technical staff of IrsiCaixa for sample processing (L. Ruiz, E. Grau, R. Ayen, L. Gomez, C. Ramirez, M. Martinez, and T. Puig).

## Authors’ contributions

SM-L and JM-P were responsible for the design and execution of the study. SM-L, JD, PC, BM, MR-R, BC, CB and JM-P were responsible for the clinical data collection, confection of the cohorts and sample collection. SM-L, LG-S, LS, VU, EW, AT and JM-P were responsible for the single-cell multiomic and statistical analysis. SM-L, VU, AG-G, NH and OJP SM-L and JM-P were responsible for metabolome and lipidome analysis. SM-L and VU were responsible for proviral and proteomic analysis. SM-L was responsible for writing the manuscript. SM-L and JM-P were responsible for the decision to submit the manuscript. LG-S, LS, VU, EW, JD, AG-G, NH, PC, BM, LR-S, MR-R, OJP, BC, CB, AT and JM-P made important critical reviews and feedback on the manuscript. The co-corresponding author had full access to all the data in the study and had final responsibility for the decision to submit for publication.

## Supplementary tables

**Table S1. Number of nuclei and reads assayed per sample (after quality control and preprocessing)**

**Table S2. Differentially accessible chromatin regions per cell type and timepoint comparison (pre- vs post-HIV, pre-HIV vs post-ART, post-HIV vs post-ART)**

**Table S3. Differentially accessible chromatin peaks (DAP) per cell type and timepoint comparison (pre- vs post-HIV, pre-HIV vs post-ART, post-HIV vs post-ART)**

**Table S4. Gene set enrichment analysis (GSEA) from DAP per cell type and timepoint**

**Table S5. Differentially expressed genes (DEG) per cell type and timepoint comparison (pre vs post HIV, preHIV vs postART, postHIV vs postART)**

**Table S6. Gene set enrichment analysis (GSEA) from DEG per cell type and timepoint**

**Table S7. HIV-positive read sequences information**

## SUPPLEMENTARY FIGURES

**Supplementary Figure 1. UMAP visualization showing the global different immune cell populations** identified at each time point (CD4, CD8, NK, proliferating NK-CD8, B, myeloid, HSPC, unknown and non-considered).

**Supplementary Figure 2. Combined differentially accessible and expressed genes.** (A) Venn diagram of the number of differentially accessible and expressed genes. (B) Number of top shared differentially accessible and expressed genes per cell type and time point comparison. (C) Hallmark gene set enrichment analysis of the shared genes per time point comparison.

**Supplementary Figure 3. Associated gene accessibility peaks and gene expression (GEX) of IP-10 (CXCL10) per time point and cell population.** In the populations with an increase in DNA accessibility and expression after HIV acquisition (A) CD4 T central memory (CM), (B) CD4 T effector memory (EM), (C) MAIT, (D) CD8 activated, (E) Proliferating NK-CD8, (F) Plasmablast and (G) conventional dendritic cells (cDC) 2.

---

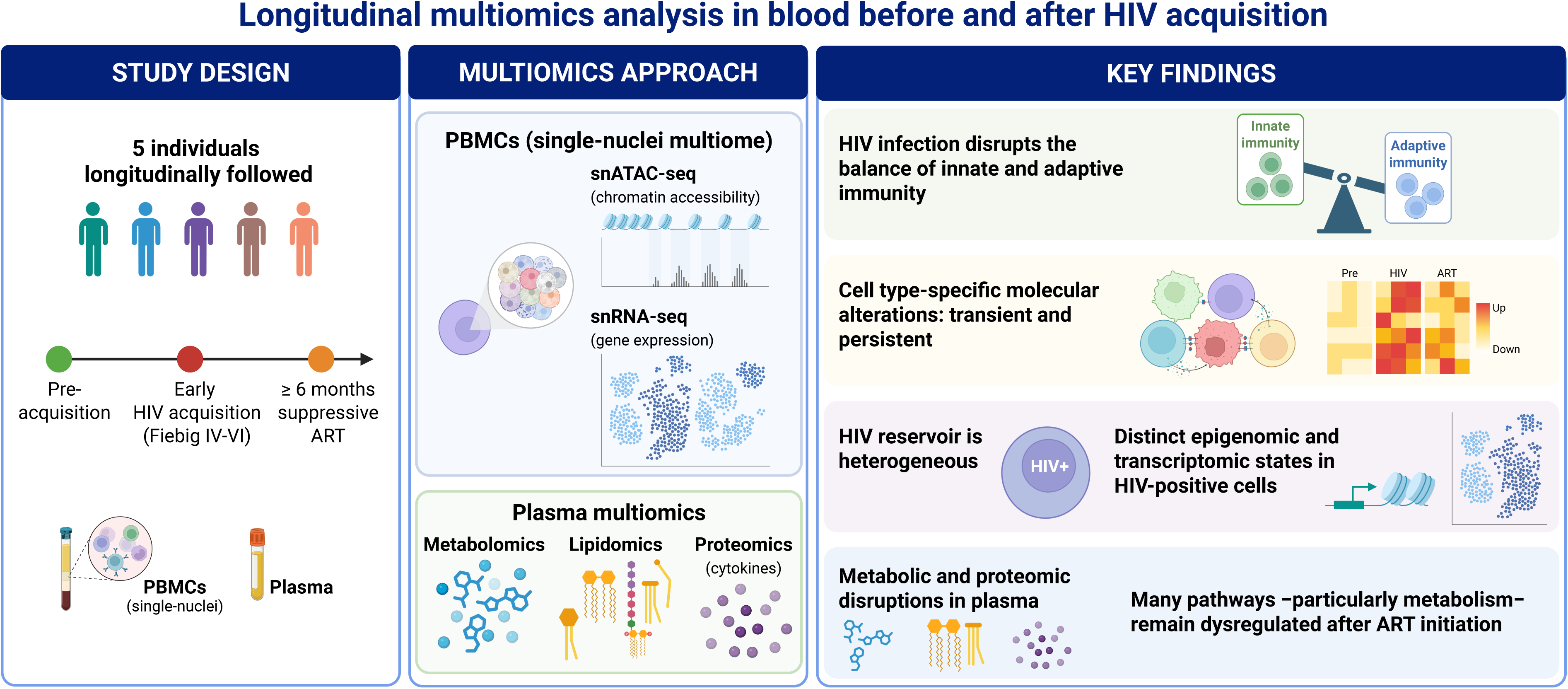

## Notes

### Competing Interest Statement

The authors have declared no competing interest.

