## Supplementary figures and images for "Longitudinal multiomics analysis in blood from individuals before and after HIV acquisition"

### Supplemental Figure 1

#### Supplementary Figure 1

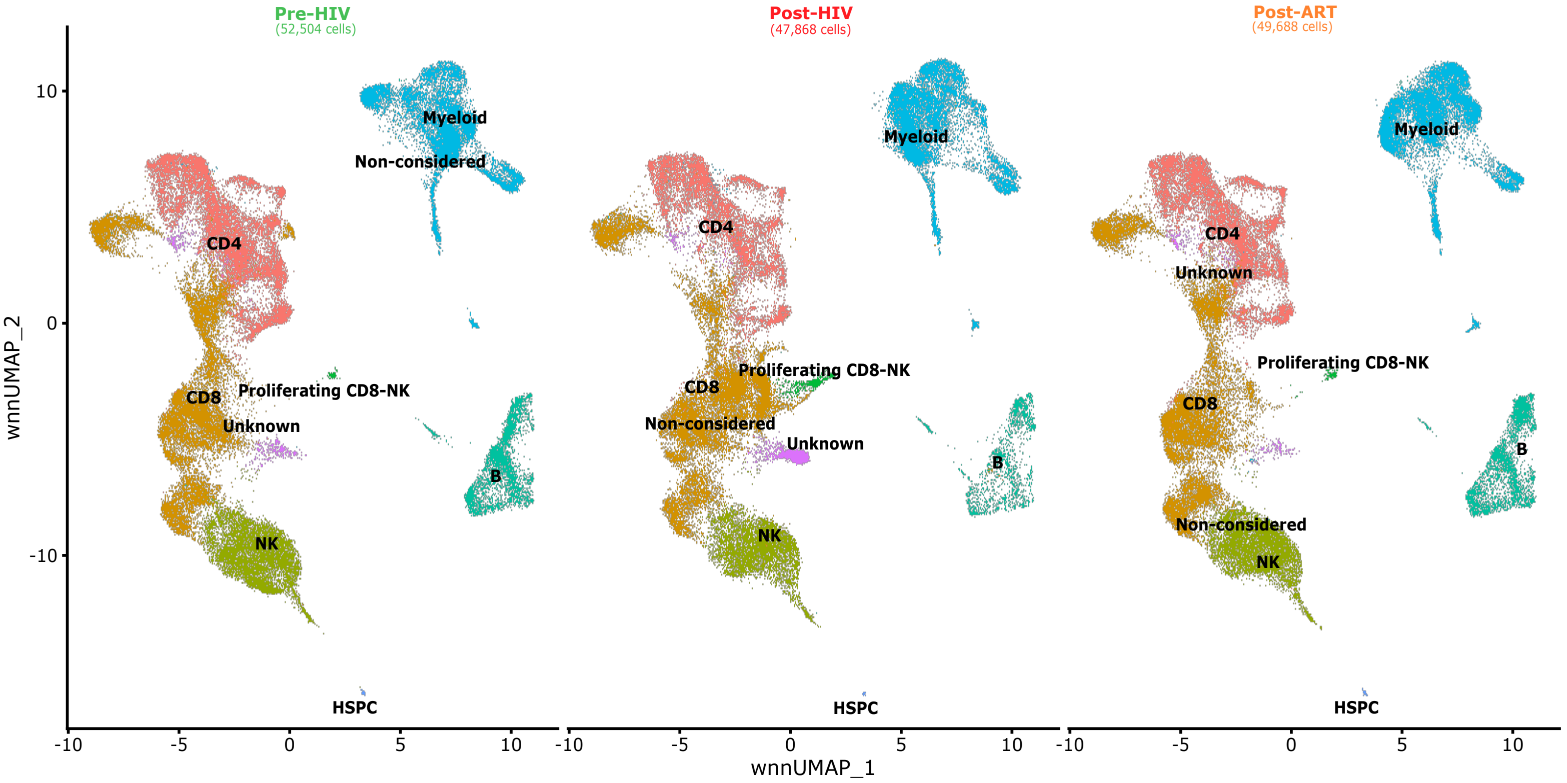

### Supplemental Figure 2

#### Supplementary figure 2

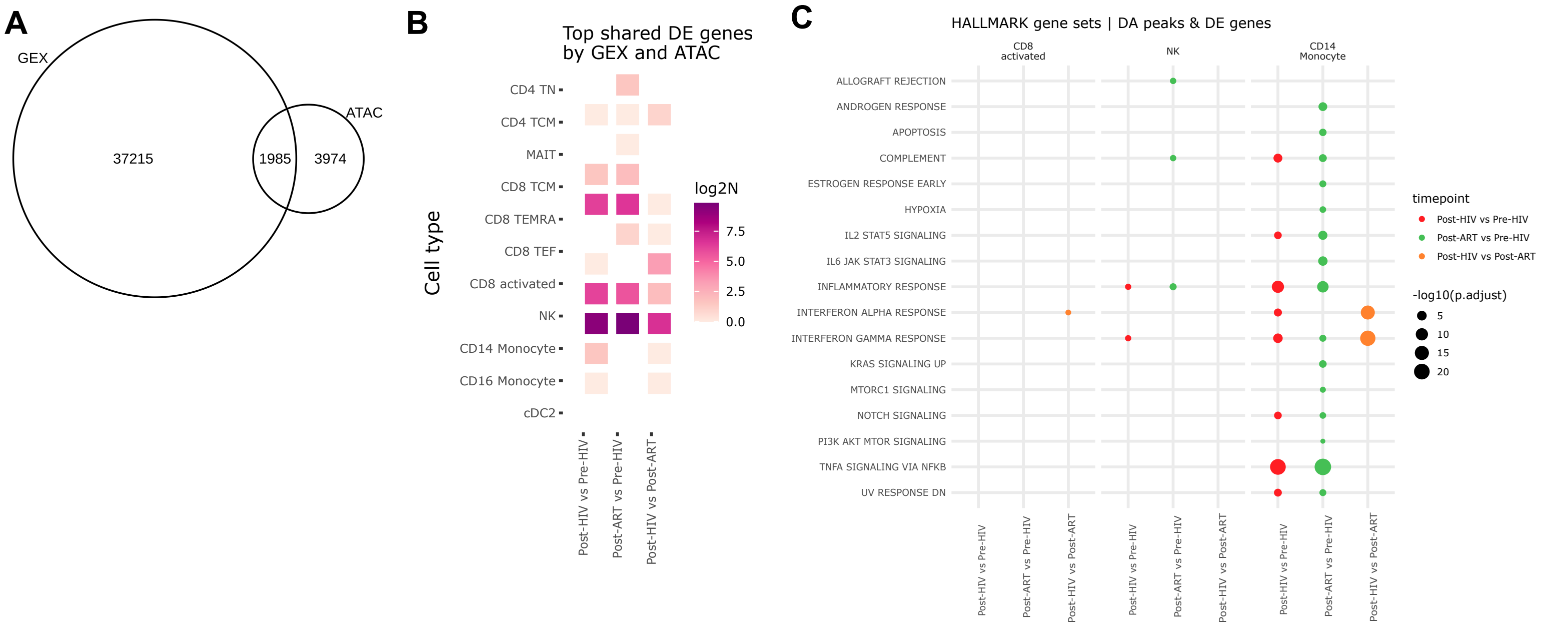

### Supplemental FIgure 3

#### Supplementary figure 3

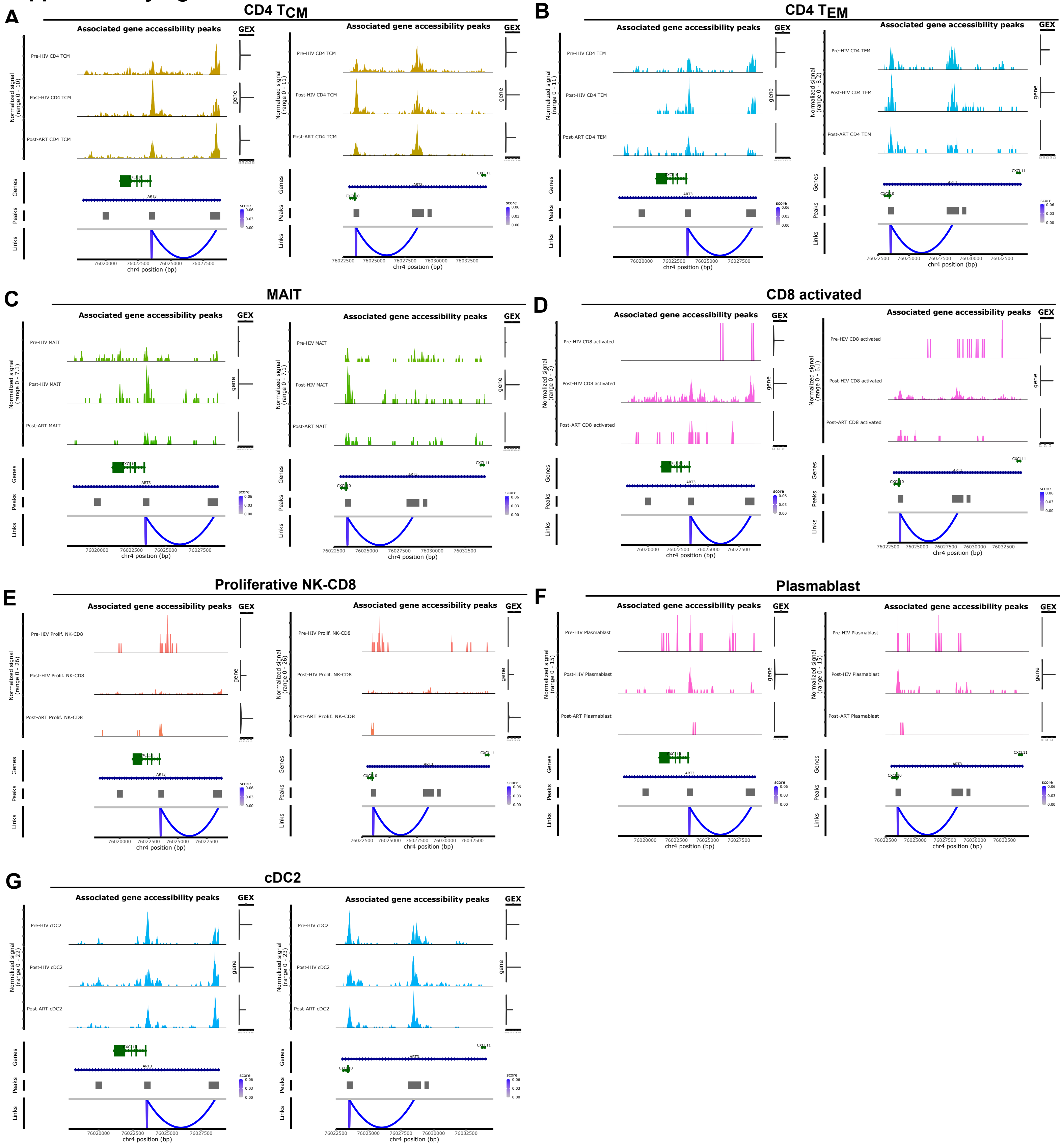
